# Structural model of PORCN illuminates disease-associated variants and drug binding sites

**DOI:** 10.1101/2021.07.19.452875

**Authors:** Jia Yu, Pei Ju Liao, Weijun Xu, Julie R. Jones, David B. Everman, Heather Flanagan-Steet, Thomas H. Keller, David M. Virshup

## Abstract

Wnt signaling is essential for normal development and is a therapeutic target in cancer. The enzyme PORCN, or porcupine, is a membrane-bound O-acyltransferase (MBOAT) that is required for the post-translational modification of all Wnts, adding an essential mono-unsaturated palmitoleic acid to a serine on the tip of Wnt hairpin 2. Inherited mutations in PORCN cause focal dermal hypoplasia, and therapeutic inhibition of PORCN slows the growth of Wnt-dependent cancers. Here, based on homology to mammalian MBOAT proteins we develop and validate a molecular structural model of PORCN. The model accommodates palmitoleoyl-CoA and Wnt hairpin 2 in two tunnels in the conserved catalytic core, shedding light on the catalytic mechanism. The model predicts how previously uncharacterized human variants of uncertain significance can alter PORCN function. Drugs including ETC-159, IWP-L6 and LGK-974 dock in the PORCN catalytic site, providing insights into PORCN pharmacologic inhibition. This structural model provides mechanistic insights into PORCN substrate recognition and catalysis as well as the inhibition of its enzymatic activity and can facilitate the development of improved inhibitors and the understanding of disease relevant PORCN mutants.

## INTRODUCTION

Wnts are secreted fatty acid-modified glycoproteins that play essential roles in both development and adult homeostasis (Loh et al., 2016; Zhong et al., 2020). Wnt ligand binding to receptors and coreceptors on the surface of target cells initiates an array of downstream signaling events, most notably β-catenin mediated transcription (Jung and Park, 2020; Nusse and Clevers, 2017; Yu and Virshup, 2014). Due to its important functions in diverse processes, the Wnt pathway is tightly regulated at multiple levels. One essential control point is the production and transport of Wnt ligands from Wnt-producing cells to target cells in the local environment.

There are 19 Wnt genes in the mammalian genome, and each requires a post-translational acylation on an essential serine positioned at the tip of disulfide-bond stabilized Wnt hairpin 2 (Bazan et al., 2012). This unique modification is catalyzed by the enzyme Porcupine (PORCN) which is present in essentially all metazoans. PORCN is a member of the family of membrane-bound *O*-acyltransferases (MBOATs) (Hofmann, 2000), and catalyzes the transfer of an monounsaturated 16:1 palmitoleate from Coenzyme A, S-(9Z)-9-hexadecenoate (PAM-CoA) to the hydroxyl of the serine residue (see model, Figure 1A) (Heuvel et al., 1993; Riggleman et al., 1990; Takada et al., 2006). Acylated Wnts are then able to transfer to the Wnt transport protein WLS and be transported to the plasma membrane (Coombs et al., 2010; Nygaard et al., 2021). The palmitoleate modification is also required for Wnt binding to its receptor Frizzled (FZD) (Hirai et al., 2019; Janda et al., 2012; Nile et al., 2017).

**Figure 1.**
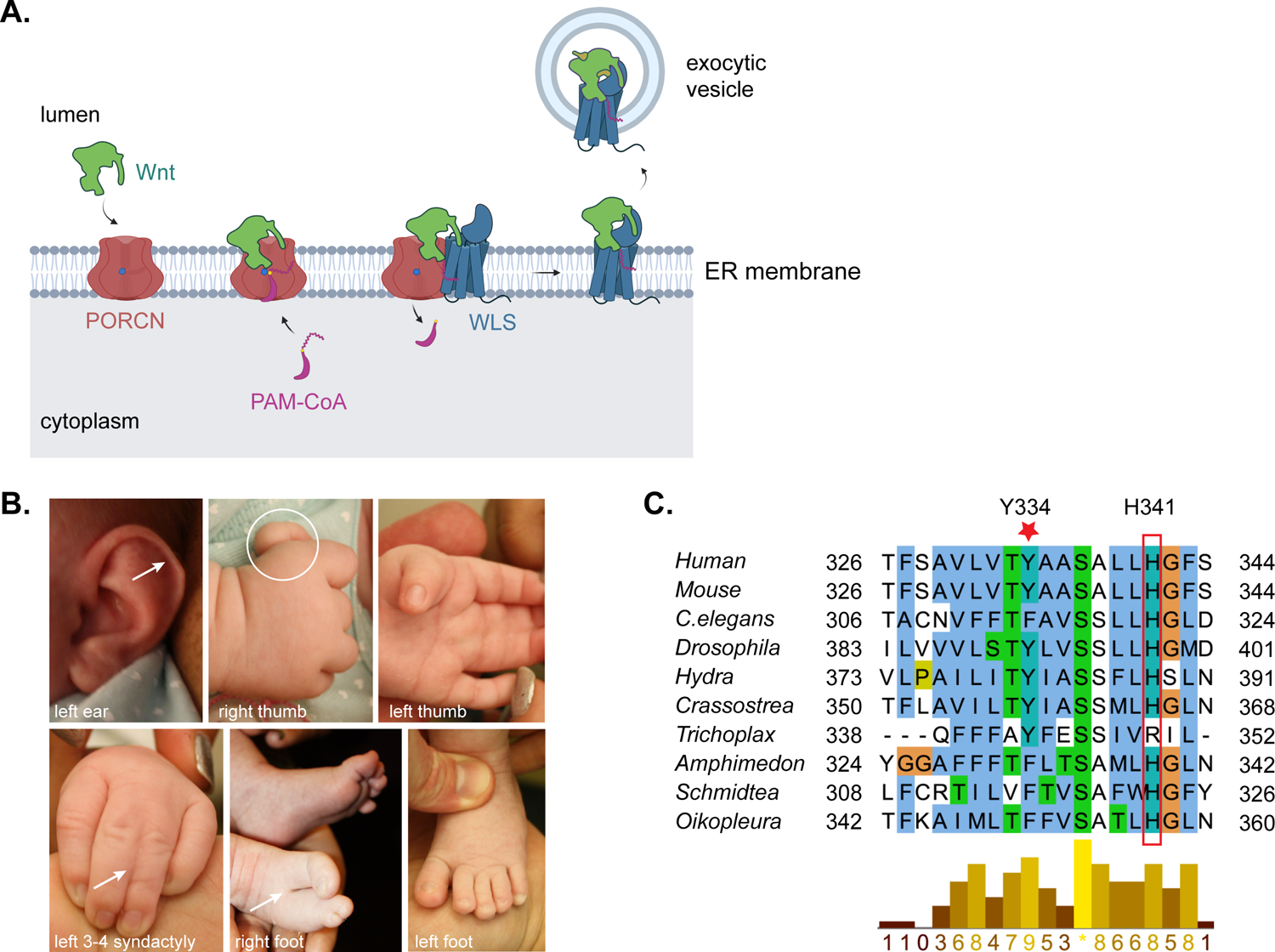
Patient clinical images and molecular findings. **(A).** PORCN is an essential Wnt O-acyltransferase. Model created with BioRender.com showing Wnt ligand (green) being acylated by PORCN (red) in the endoplasmic reticulum (ER) membrane using the cofactor palmitoleoyl-CoA (Pam-CoA, purple). After acylation, Wnt is transferred to the carrier protein WLS (blue) and subsequently enters the secretory pathway. **(B).** Multiple physical abnormalities including thinning of the left ear helix (top left, arrowhead), hypoplasia of both thumbs, left finger 3-4 syndactyly (bottom left, arrowhead), right foot with a central cleft, absence of toes 2 and 3, and syndactyly of toes 4 and 5 (bottom middle, arrowhead) and a normal left foot (bottom right) in a patient with suspected focal dermal hypoplasia. **(C).** Multiple sequence alignment of metazoan PORCN proteins, illustrating the conserved Y334 residue (red star). Some sequences were obtained by iterative PsiBLAST searches in the genomes of Cnidaria, Placozoa, and Porifera genomes. The sequences used here for alignment are from *Drosophila melanogaster* (NP_476890.1) belonging to phylum arthropods. *C.elegans* (NP_509054.1) which belongs to phylum Nematoda. Hydra (Hydra vulgaris, XP_012563063.1) is Cnidaria. Crassostrea (*Crassostrea virginica*, XP_022296962.1) belongs to the phylum Mollusca. Trichoplax (Trichoplax sp. H2, RDD44615.1) is in the phylum Placazoa. The sponge Amphimedon (*Amphimedon queenslandica*, XP_003387071.1) is in the phylum Porifera. Schmidtea (*Schmidtea mediterranea*, ABW97518.1) is planaria, in the phylum Platyhelminthes. Oikopleura (*Oikopleura dioica*, CBY33632.1) is a sea squirt in the phylum Chordata. The human PORCN (isoform D, NP_982301.1) and mouse (isoform D, NP_076127.1) sequences are included at the top as references. Alignment and conservation scores were determined using Clustal Omega online server (https://www.ebi.ac.uk/Tools/msa/clustalo/) and Jalview with the ClustalX color scheme. The catalytic site H341 is highlighted by a red box.

Because Wnt signaling drives an important subset of cancers and PORCN function is essential for Wnt activity, a growing number of small molecules that potently inhibit the catalytic activity of PORCN have been developed (Shah et al., 2021; Zhong and Virshup, 2020). These drugs block the palmitoleation of Wnts and inhibit the growth of Wnt-addicted cancers. Several of these PORCN inhibitors are in human clinical trials. How and where these drugs interact with the PORCN structure to block its activity is not known, although pharmacophore models have been developed to identify critical features of the drugs (Poulsen et al., 2015).

Wnt signaling is also critically important during embryonic development (Loh et al., 2016). Loss of function mutations of PORCN and WLS that block the acylation and delivery of Wnts cause early embryonic lethality in mouse knockout models (Barrott et al., 2011; Biechele et al., 2011; Fu et al., 2009). Multiple congenital malformation syndromes are due to partial loss of function mutations in Wnt production. Among those, diverse germ line mutations in *PORCN* are associated with Goltz syndrome, an X-linked genetic disorder also known as focal dermal hypoplasia (FDH; OMIM 305600) (Bostwick et al., 2016; Goltz, 1992; Grzeschik et al., 2007; Wang et al., 2007). This rare disorder affects the development of ectodermal and mesodermal tissues, and affected individuals have variable defects in limbs, skin, and bone as well as other abnormalities in internal organs. FDH is usually found in heterozygous females where the severity of the disease depends on the pattern of X-inactivation or the presence of somatic mosaicism. Males may also have FDH, usually in the setting of a mosaic mutation, although germline mutations in *PORCN* have been identified in hemizygous males with colobomata, microphthalmia, anophthalmia or other findings of FDH (Brady et al., 2015; Madan et al., 2017; Wawrocka et al., 2021).

Overall, more than 170 unique genomic variants in the *PORCN* locus linked to FDH have been listed in the Global Variome shared LOVD database (https://databases.lovd.nl/shared/genes/PORCN). Many of these are classified as Variants of Uncertain Significance (VUS) because they are missense mutations located outside of the known functional features. Determining if these variants are deleterious can be facilitated by analysis of comparative evolutionary and where available, structural features and computationally can be estimated by online tools such as PolyPhen-2 (Adzhubei et al., 2010), Missense3D (Khanna et al., 2021), and PROVEAN (Choi and Chan, 2015). Laboratory-based assessment of PORCN function, an approach we and others have used previously, is significantly more laborious (Proffitt and Virshup, 2012; Rios-Esteves et al., 2014). Prioritizing which mutants to test in cell-based assays would be facilitated by knowledge of where the variants lie in the PORCN structure. While the PORCN structure has not been solved, the structures of a bacterial and two mammalian MBOAT proteins were recently determined. They can provide a homology template to model PORCN structure and hence guide a more refined assessment of the consequences of missense mutations and the location of PORCN inhibitor binding sites.

Here, taking advantage of advances in structural prediction algorithms, we developed a high-quality structural homology model of human PORCN (Sánchez and Šali, 1997; Vyas et al., 2012; White, 2009). Based on recently published structures of human MBOAT proteins (Long et al., 2020; Qian et al., 2020; Sui et al., 2020; Wang et al., 2020), this PORCN model is consistent with experimentally determined topology and functional residues, explains the effects of multiple FDH mutations, and predicts the consequences of new human variants of uncertain significance.

Molecular docking of WNT8A and PAM-CoA in the structure positions the PORCN catalytic histidine 341 adjacent to the thioester bond and the serine in Wnt hairpin 2, providing a model for Wnt acylation. Multiple PORCN inhibitors including ETC-159, LGK974, and IWP-L6 can be docked into the PORCN active site in a manner consistent with an established pharmacophore model (Poulsen et al., 2015). This structural model provides mechanistic insights into PORCN substrate recognition and catalysis as well as the pharmacologic inhibition of its enzymatic activity and can facilitate the development of improved inhibitors and the rationalization of disease-relevant human PORCN variants.

## MATERIALS AND METHODS

### Ethical considerations

The patient was seen at the Greenwood Genetic Center (GGC), Greenwood, South Carolina, for genetic evaluation related to congenital limb anomalies. Informed consent was signed by the parent of the proband prior to participation in the research study. Consent included permission to publish clinical information and photographs. All procedures employed were in accordance with study parameters approved by the Institutional Review Board, and compliant with practices at the GGC.

### PORCN functional assays

3xHA tagged mouse PORCN-D encoding plasmids were described previously (Proffitt and Virshup, 2012; Tanaka et al., 2000). Site-directed mutagenesis was performed to generate PORCN mutants, and the sequences were verified by Sanger sequencing. PORCN-knockout HT1080 cells were generated and characterized previously (Proffitt and Virshup, 2012). To monitor Wnt secretion into the culture medium, cells were grown in 12-well plates and transfected with 500 ng of WNT3A expression plasmid, along with 50 ng and 100 ng of expression plasmids encoding 3xHA-tagged wildtype or mutant mouse PORCN-D as indicated using Lipofectamine 2000 (Thermo Fisher). 24 hours after transfection, the medium was changed to low serum (2% FBS) at half volume (0.5 ml). For Wnt secretion experiments, medium was collected after another 24 hours and subjected to western blotting. For SuperTOPFlash assays, experiments were performed in 24-well plates, transfecting 200 ng of SuperTOPFlash plasmid together with 50 ng of WNT3A, 100 ng of mCherry, and up to 20 ng of PORCN expression plasmids. 24 hours after transfection, luciferase activity was quantitated as previously reported (Yu et al., 2020). If Wnt inhibitors were used, the cells were treated with the indicated inhibitors beginning 6 hours after transfection and incubated for another 18 hours before harvesting for the reporter assay.

### Homology modelling of PORCN structure

Homology modeling was performed using the MODELLER (Webb and Sali, 2016) program from trRosetta (Yang et al., 2020) and (PS)^2^ (Huang et al., 2015) by query-template alignment. Two recently published human MBOAT proteins structures, DGAT1 (PDB 6VP0) and ACAT1 (PDB 6P2P), were chosen as templates to guide the modelling with the human PORCN amino acid sequence (Unipro:Q9H237-1/NCBI Q9H237.2, also known as splice isoform D, 461 amino acids), using the T-Coffee homology extension (PSI-coffee) algorithm (Tommaso et al., 2011). To refine the PORCN simulation, I-Tasser (Yang and Zhang, 2015a) was used by assigning the template generated from trRosetta and (PS)^2^ with specified secondary structure for the C-terminal 41 amino acids (Val421 to Gly461). Lastly, the simulated structure was refined by GalaxyRefine2 from GalaxyWEB (Afgan et al., 2018; Boekel et al., 2015) to generate the final predicted structure.

Protein tunnels were identified and assessed using Caver 3.03 (Pavelka et al., 2015) as a Pymol Plugin using a probe radius of 2 Å, shell radius of 4 Å, and a shell depth of 3 Å. The clustering threshold was set at 3.5. The starting point was defined as the point between His341, Asn306, and Phe257.

### Molecular Docking of Palmitoleoyl-CoA, WNT8A and PORCN inhibitors

System preparation and receptor-ligand docking calculations were performed using the Schrödinger Suite package (version 2020–4), using default parameters unless otherwise noted. The homology model of human PORCN as a receptor was first prepared using the Protein Preparation Wizard. The atom and bond types were corrected and the protonation states of ionizable species adjusted to pH 7.4 by Epik. During receptor grid generation, the ligand to be docked was confined to an enclosing box centered on the middle of the tunnel, using Ser262, Glu300, Asn315 and His341 to set the box center with an external box of 36 Å for PAM-CoA docking, and a ligand-sized box centered on Glu293, Val302 and His409 for inhibitor docking. The PAM-CoA structure and PORCN inhibitors were generated from PubChem, prepared using the LigPrep module to convert the 2D sdf format into 3D molecular structures, and docked to the orthosteric site of PORCN with the Glide program (Friesner et al., 2006) using the standard precision (SP) mode. The best binding pose predicted for human PORCN-PAM-CoA complex and PORCN-inhibitors docking was saved for further analysis for molecular interactions.

To predict protein-protein interactions, the structure of WNT8A was extracted from the WLS/WNT8A structure (PDB 7KC4) (Nygaard et al., 2021) and the ClusPro 2.0 Server (Kozakov et al., 2017) was used to predict its interaction with PORCN. We used the hydrophobic-favored potential in the server. A near-native state of protein conformations was chosen during protein-protein docking.

## RESULTS

### Whole exome sequencing identified a novel heterozygous variant in the *PORCN* gene in a patient with features consistent with Focal Dermal Hypoplasia

The proband initially presented to the Greenwood Genetics Center clinic at 6 weeks of age for evaluation of congenital limb anomalies (Figure 1B). She also had a history of mild right branch peripheral pulmonary artery stenosis. Her physical exam was notable for abnormalities of both hands and the right foot. On the left hand, there was a hypoplastic left thumb containing a nail, cutaneous syndactyly of fingers 3-4 extending to the bases of the distal phalanges, and mild medial clinodactyly of the 5th finger. On the right hand, there was a hypoplastic thumb containing a nail, mild radial deviation of the 2nd finger with a slight crease between the bases of fingers 2-3, and a suggestion of syndactyly at the bases of fingers 3-4. The right foot had a medially incurved hallux, absence of toes 2 and 3 with a central cleft, and almost complete syndactyly of toes 4 and 5 with separate nails. The right leg also appeared slightly shorter than the left, and there was mild elevation of the external genitalia and lower buttock on the right side as compared to the left. Other minor atypical features seen on physical exam included thinning of the left ear helix, mild asymmetry of the nasal alae, retrognathia, and inverted nipples. The hair was thinner on the right parietal scalp as compared to the left, and no skin abnormalities were noted. A previous eye exam had shown mildly anomalous irises (without colobomas), a minimal congenital nuclear cataract in the right eye, and bilateral astigmatism, while subsequent eye exams showed the additional findings of bilateral nasolacrimal duct obstruction (ultimately requiring dilatation) and small optic nerves but no evidence of a cataract.

A blood chromosome analysis and whole genome chromosomal microarray were normal. Whole exome sequencing was performed as previously described (Spellicy et al., 2019), with an initial focused analysis of the *TP63* and *SALL4* genes that were normal. Additional analysis of the remaining exome sequencing data revealed a novel heterozygous p.Tyr334Cys (c.1001A>G) variant of uncertain significance in the *PORCN* gene located on the X chromosome. Tyr334 is highly conserved in PORCN across diverse species and is only seven residues away from the catalytic His341 (Figure 1C) (Hofmann, 2000), suggesting the mutation could be pathogenic. Targeted testing of the patient’s mother showed her peripheral blood lymphocytes to be homozygous normal (c.1001A), and the patient’s father was unavailable for testing. X-inactivation analysis on the patient was deemed uninformative. Together with the proband’s clinical features, her variant in *PORCN* was felt to be consistent with a diagnosis of focal dermal hypoplasia (FDH), an X-linked dominant condition in which Split-hand/foot malformation and other anomalies can occur. Considering this diagnosis, additional evaluations were pursued to screen for other potential findings of FDH. An abdominal ultrasound showed no evidence of renal or diaphragmatic anomalies. An audiologic evaluation was normal, and an ENT evaluation showed no laryngeal papillomas.

### Homology modeling of human PORCN structure

This case adds to the many variants of uncertain significance (VUS) in *PORCN* that have been reported in various databases. Prediction of phenotype from genotype is an inexact science and testing every individual VUS in cell-based assays is not widely available. We speculated that a structural model of PORCN would aid in predicting the phenotypic consequences of the VUS associated with focal dermal hypoplasia. PORCN is a multipass-transmembrane protein that localizes in the endoplasmic reticulum (ER), and while its topology has been experimentally established with protein tagging (Galli et al., 2021; Lee et al., 2019), there is currently no experimentally-derived structure. The three-dimensional structures of a bacterial and more recently several human MBOAT family members have been determined that provided mechanistic insights into their enzymatic functions (Jiang et al., 2021; Long et al., 2020; Ma et al., 2018; Qian et al., 2020; Sui et al., 2020; Wang et al., 2020). In addition, recent advances in modeling are increasingly capable of producing accurate structures (Yang et al., 2020). We therefore undertook a comparative protein structure modeling approach leveraging off these new human MBOAT structures.

To select the optimum template for PORCN modeling, we compared sequence alignments of PORCN with the MBOAT proteins with solved structures, including DltB, ACAT1 and DGAT1 (Supplemental Figure 1). As a first approach to generate a PORCN structural model, human PORCN sequence was uploaded to several platforms including Phyre (Kelley et al., 2015), Rosetta (DiMaio et al., 2011), RaptorX (Källberg et al., 2014), Swiss-model (Waterhouse et al., 2018) and I-Tasser (Yang and Zhang, 2015a, 2015b). These programs selected the bacterial MBOAT DltB (Ma et al., 2018) as the homology template for PORCN due to the highest score of identity (14%). However, none of the simulated structures from these platforms could be docked with PAM-CoA successfully due to the small tunnel size.

We next used MODELLER (Webb and Sali, 2016) to generate 3D structure models for human PORCN by using the MBOAT structures diacylglycerol O-acyltransferase 1 (DGAT1) (Wang et al., 2020) and acyl-coenzyme A:cholesterol acyltransferase (ACAT1) (Long et al., 2020; Qian et al., 2020) as templates. The C-terminal 40 amino acids of PORCN lack similarity to the other MBOATs and were therefore refined using I-TASSER, which is ideal for modeling sequences of low homology via thread methodology (see Methods) (Yang and Zhang, 2015a). The homology structure based on DGAT1 (Figure 2) was deemed to be of highest quality, with 92.2% of residues in favored and another 6.1% in allowed Ramachandran regions. In addition, the structure could readily be docked with PAM-CoA and WNT8A (discussed below) and was therefore further evaluated.

**Figure 2.**
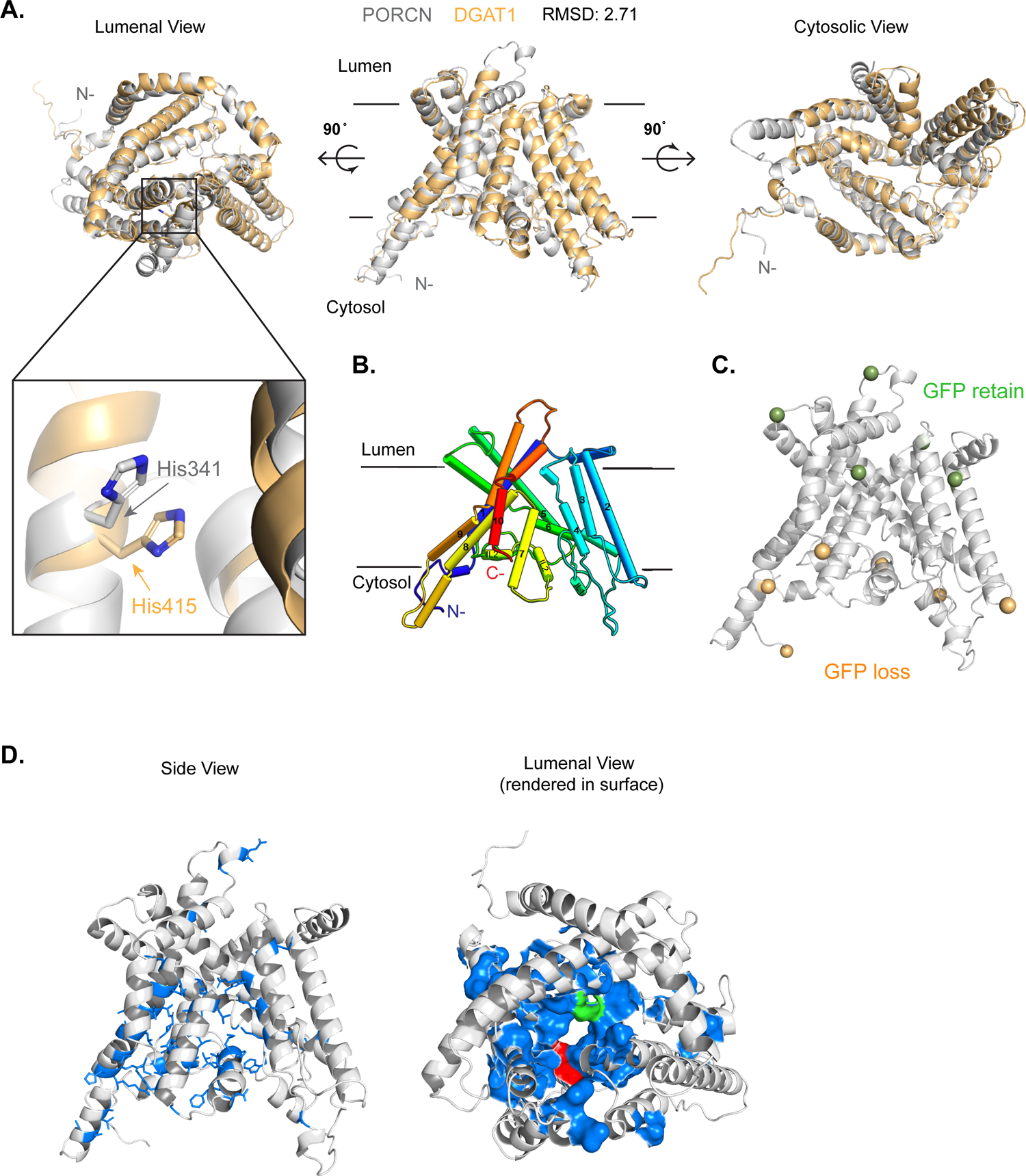
PORCN structure homology modeling. **(A).** Three views of the PORCN homology model (in grey) and DGAT1 (PDB 6VP0) structure (in light orange) superimposed. The RMSD of 2.71 Å is calculated on the Cα backbone. The lumenal side is where the substrate Wnt protein enters, and the cytosolic side is where the donor palmitoleoyl-CoA enters. Inset shows that the catalytic residues His341 (for PORCN) and His415 (for DGAT1) are in close proximity. **(B).** Topology of the PORCN homology model, highlighting the ten TM helices in rainbow color from the N (blue) to the C (red) terminus. **(C).** The PORCN model recapitulates the results of the GFP protection assay of (Lee et al., 2019). The spheres in the model indicate where GFP was inserted; green indicates protection from, while orange indicates sensitivity to protease digestion. Protease-resistant sites reside on the ER lumenal side, while protease-sensitive sites are on the cytosolic side. **(D).** Co-evolving residues reside in the PORCN catalytic core. A co-evolution score was determined from alignment of 468 sequences, and the position of the top co-evolving residues (referred to as a coupling block) are indicated in blue sticks in the side view of the structure. Co-evolving residues are also lining the predicted catalytic core in the membrane (view from ER lumen rendered in surface view in the right panel). Red residue indicates His341, and green residue indicates Asn306.

Overall consistent with the experimental data of (Lee et al., 2019) and (Galli et al., 2021), the predicted PORCN model has 10 transmembrane helices with both the N- and C-termini in the cytoplasm. Superimposition of the PORCN structure with DGAT1 showed a root mean square deviation (RMSD) of only 2.71 Å (Cα backbone) (Figures 2A and 2B). The PORCN structure also has similar TM structure and folds as DltB and ACAT1 (Supplemental Figure 2A). The catalytic histidine (His341) of PORCN is located in the middle of a tunnel, in a similar position as the catalytic residue His415 in DGAT1 (Figure 2A, inset). The model with the topology reported by the fluorescence protease protection studies of Lee et al. is shown in Figure 2C (Lee et al., 2019).

As an independent approach to evaluate the quality of the model, we used it to map two features of PORCN evolution: amino acid conservation, and amino acid co-evolution. Residues critical for the function of a protein tend to be conserved, and indeed, mapping sequence conservation determined with ConSurf (Ashkenazy et al., 2016) onto the structure of PORCN showed clustering of highly conserved residues in both the catalytic core and the presumed Wnt and PAM-CoA binding pockets (Supplemental Figure 2B). Additional useful information can be deduced from how amino acid networks co-evolve. This is based on the principle first recognized by Darwin, that biological structures (or in the case of proteins, residues) that are functionally linked tend to evolve in tandem (reviewed in (Göbel et al., 1994; Juan et al., 2013; Ovchinnikov et al., 2015)). Given the large number of sequences available, we used computational methods to visualize PORCN co-evolving blocks. Analysis of 468 PORCN amino acid sequences from species as diverse as sponge and human using the LERI server identified several co-evolving blocks (Cheung et al., 2021). The most highly co-evolving residues, indicated by the blue block in Supplemental Figure 2C (and see example in Supplemental Figure 2D), when mapped on to the PORCN structure (Figure 2D) highlight the inner catalytic core of the enzyme, supporting the functional accuracy of the PORCN structure prediction.

### PORCN transmembrane tunnel analysis

PORCN is an O-acyl transferase that preferentially catalyzes the transfer of the cis-Δ9 unsaturated palmitoleate (C16:1) to a conserved serine on hairpin 2 of Wnts. Based on homology to ACAT1 and DGAT1, we examined the PORCN structure for the presence of tunnels that might accommodate these ligands. Indeed, CAVER software highlighted two tunnels, denoted 1 and 2, in the TM domain that open to the ER lumenal and cytoplasmic sides of the membrane respectively (Figure 3A). Supporting their relevance, the residues in PORCN that are most highly conserved (Ashkenazy et al., 2016) form the walls of these contiguous structural tunnels 1 and 2 (Figure 3A).

**Figure 3.**
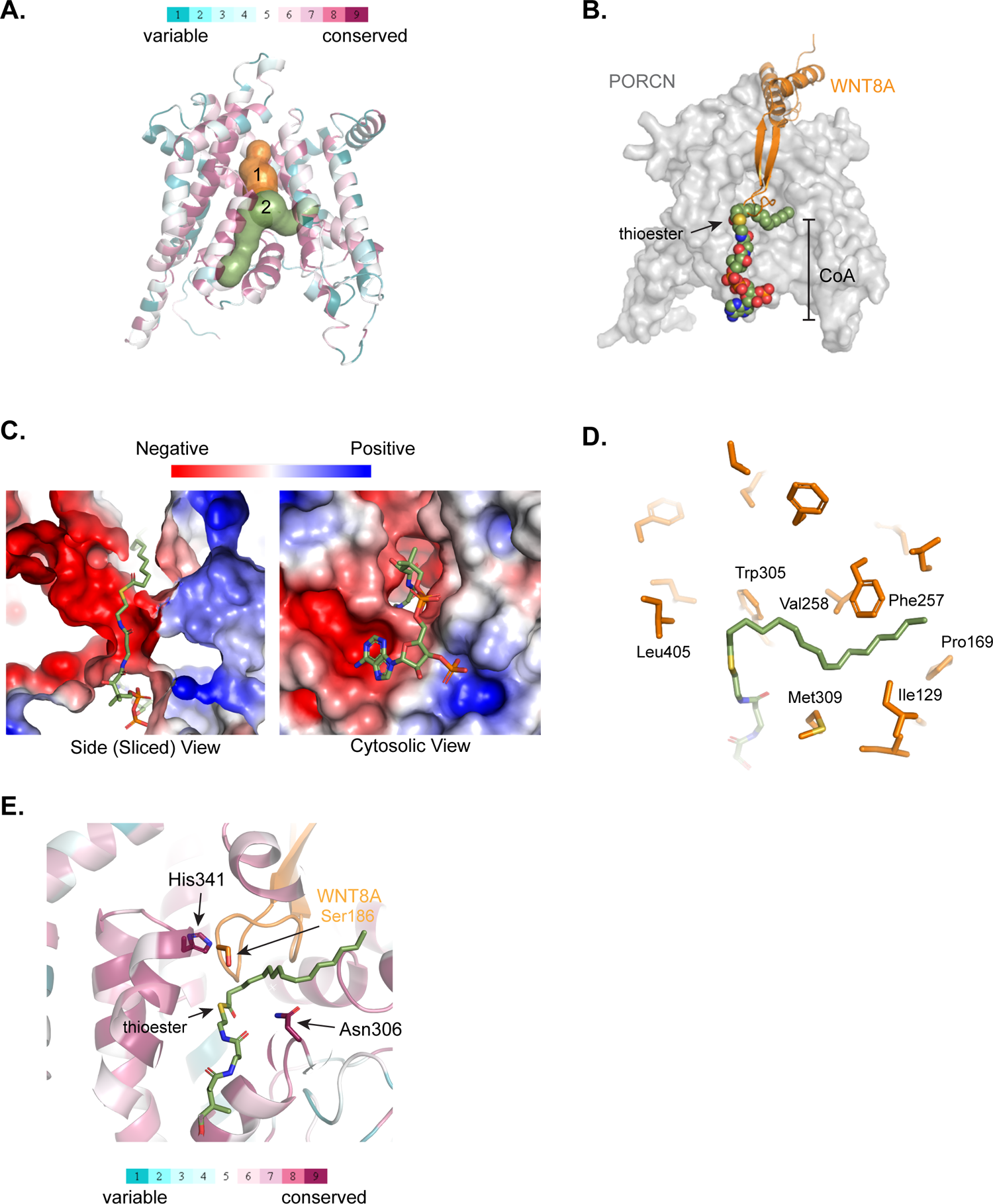
PORCN model predicts Wnt hairpin 2: palmitoleoyl-coenzyme interaction. **(A).** Conservation map and tunnel prediction for PORCN. Two tunnels were predicted using CAVER, with the orange tunnel 1 open to the ER lumen (top) and the green tunnel 2 open to the cytosol (bottom). The model is colored according to sequence conservation, highlighting the high conservation of residues surrounding the two cavities. The green tunnel 2 has a kink in the middle that could accommodate a cis-palmitoleate. **(B).** PAM-CoA and WNT8A were sequentially docked onto the PORCN structure. Only WNT8A hairpin 2 (orange) is shown for clarity. PORCN is represented in semi-transparent surfaces and PAM-CoA is represented by spheres. **(C).** Slice of the molecular electrostatic surface model showing the interaction of the CoA moiety aligned with complementary charged surfaces in the cytosolic cavity of the PORCN model. The electrostatic surface was calculated using the APBS function of Pymol. **(D).** Molecular interactions of the cis-palmitoleate in tunnel 2. The acyl chain of the PAM-CoA is surrounded by an extended hydrophobic cavity of the PORCN model. **(E).** The conserved catalytic core of PORCN positions with PAM-CoA and Wnt hairpin 2 for catalysis. The essential catalytic His341 is positioned near the hydroxyl group of Ser186 of WNT8A, which in turn lies above the thioester bond. PAM-CoA is represented as a green stick figure with the thioester bond in the center (sulfur in yellow), and the position of Ser186 of WNT8A is indicated. The position of the highly conserved residue Asn306 is also indicated.

### Molecular docking of substrates provides mechanistic insights into Wnt palmitoleation

The structure and cavity analysis suggests that the Wnt acylation reaction proceeds via PAM-CoA binding to tunnel 2 via the cytoplasmic face of PORCN, and the acceptor hairpin 2 of Wnt entering the catalytic PORCN core via tunnel 1 from the ER lumen. Prior biochemical analysis suggests that PORCN His341 serves as a base catalyst to facilitate the transfer of the palmitoleoyl group from CoA to the serine of Wnt hairpin 2 (Lee et al., 2019; Nile and Hannoush, 2016; Proffitt and Virshup, 2012; Rios-Esteves et al., 2014). To examine whether our model can provide a mechanistic rationale for the molecular events during Wnt signaling, we docked PAM-CoA into the homology model of PORCN (Figure 3B-E and Supplemental Figure 3B). The CoA moiety anchors in a pocket on the cytoplasmic face (Figure 3C), similar to how it interacts with ACAT1 (Qian et al., 2020) and DGAT1 (Sui et al., 2020; Wang et al., 2020). The PAM-CoA thioester bond lies at the base of tunnel 1 (Figure 3B). The unsaturated fatty acid chain extends into tunnel 2, lined by conserved hydrophobic residues (Figure 3D-3E, Supplemental Figure 3C). This tunnel has a kink lined by PORCN Asn306 and Met309 that accommodates the *cis* double bond (Supplemental Figure 3D). We note that Asn306 is invariant in PORCN and has previously been proposed to assist in catalyzing the transesterification reaction. However, mutation of Asn306 has no effect on the catalytic activity of PORCN in cell-based assays (Proffitt and Virshup, 2012; Rios-Esteves et al., 2014) but markedly decreased activity in a purified system where only PAM-CoA was available as a substrate (Lee et al., 2019). The structural model thus suggests Asn306 could enforce the use of cis-double bond-containing fatty acid substrates (Tuladhar et al., 2019).

We next docked WNT8A (extracted from the WNT8A/WLS structure, PDB 7KC4) (Nygaard et al., 2021) into the PORCN model using the ClusPro 2.0 Server (Figure 3B and Supplemental Figure 3B). Supporting our hypothesis, the predicted complex shows WNT8A docking into tunnel 1 from the lumenal face of PORCN, with the hairpin 2 loop inserted deeply into the tunnel (Figure 3B). Interestingly, WNT3 from the WNT3: FZD complex (PDB 6AHY) did not dock hairpin 2 as deeply into the tunnel due to interference from Wnt hairpin1. Unlike WNT8A in complex with WLS, in the WNT3: FZD complex hairpin 1 is tightly apposed to hairpin 2 (Hirai et al., 2019; Nygaard et al., 2021). When the modeled complex of WNT8A: PORCN was combined with the PORCN-PAM-CoA model, the hairpin 2 loop of WNT8A is in close contact with the thioester of the PAM-CoA (Figure 3E). During Wnt acylation, PORCN catalyzes the transfer of the palmitoleoyl moiety from CoA to the conserved serine residue on Wnt hairpin 2 (Ser186 of WNT8A). In the modeled ternary complex, Ser186 is positioned 4.3 Å from the thioester bond of the PAM-CoA, and the conserved catalytic residue His341 from PORCN is well-positioned to activate the hydroxyl group of Ser186 from WNT8A for nucleophilic attack on the thioester group of the substrate (Figure 3E). Asn306, another highly conserved residue in PORCN, while close to the PAM-CoA, has a less clear role in catalysis (Lee et al., 2019; Proffitt and Virshup, 2012; Rios-Esteves et al., 2014). The structural prediction of PORCN docked with PAM-CoA and WNT8A provides a coherent and useful model to understand Wnt acylation.

Resh and co-workers reported an extensive mutational analysis of PORCN (Rios-Esteves et al., 2014). We examined where mutants that affected PORCN activity in their assays were in our predicted structure. Trp305 interacts with the thioester of PAM-CoA, Tyr316 stabilizes the interaction of IL2 with TM4, Tyr334 is altered in our patient and is discussed below, Ser337 interacts with the CoA moiety, and Leu340 is adjacent to the catalytic H341 and contacts Wnt hairpin 2 (Supplemental Figure 3E). Thus, these engineered loss-of-function mutants can also largely be explained by the predicted PORCN complexes.

### Functional analyses of the p.Tyr334Cys PORCN variant

The above analyses demonstrate that the PORCN homology model is predictive of function. Next, we assessed whether the model can be used to predict the consequences of PORCN sequence variants found in patients (Figure 4A). First, we asked if the Y334C variant had damaging consequences. Tyr334 is located on TM helix 7, the same helix as the catalytic residue His341, and appears to stabilize the PORCN catalytic core by interacting with multiple hydrophobic residues in both TM 8 (Val350, Leu354, Ile357) and on the membrane side of intracellular loop 2 (Val301 and Val302) (Supplemental Figure 4A). Consistent with this, the Missense3D server (http://missense3d.bc.ic.ac.uk/missense3d/) (Khanna et al., 2021) predicts the Y334C substitution significantly alters the internal cavity of the protein. To experimentally test the stability and Wnt/β-catenin signaling activity of this PORCN variant, we used our previously described *PORCN* knockout HT1080 cell line. This is a mesoderm-derived human male fibrosarcoma cell line where an introduced frameshift mutation inactivates the *PORCN* gene so the cells neither palmitoleate nor secrete Wnt ligands (Proffitt and Virshup, 2012). Ectopic expression of PORCN rescues Wnt secretion and can therefore be used to test the activity of various PORCN mutants. Consistent with the structural and computational prediction, PORCN Y334C has decreased protein expression in cell lysates compared to wildtype PORCN (Figure 4B). In the presence of PORCN Y334C, significantly less WNT3A was secreted into the culture medium compared to the control (Figure 4B) and the mutant also had decreased activity in a functional Wnt/β-catenin signaling assay (Figure 4C). This instability is not due to the presence of the cysteine residue, as Rios-Esteves et al, (Rios-Esteves et al., 2014) reported that PORCN Y334A also had decreased stability and decreased activity. These findings demonstrate that VUS PORCN Y334C is a deleterious mutation responsible for the patient’s disease and support the value of the PORCN model to predict functional consequences.

**Figure 4.**
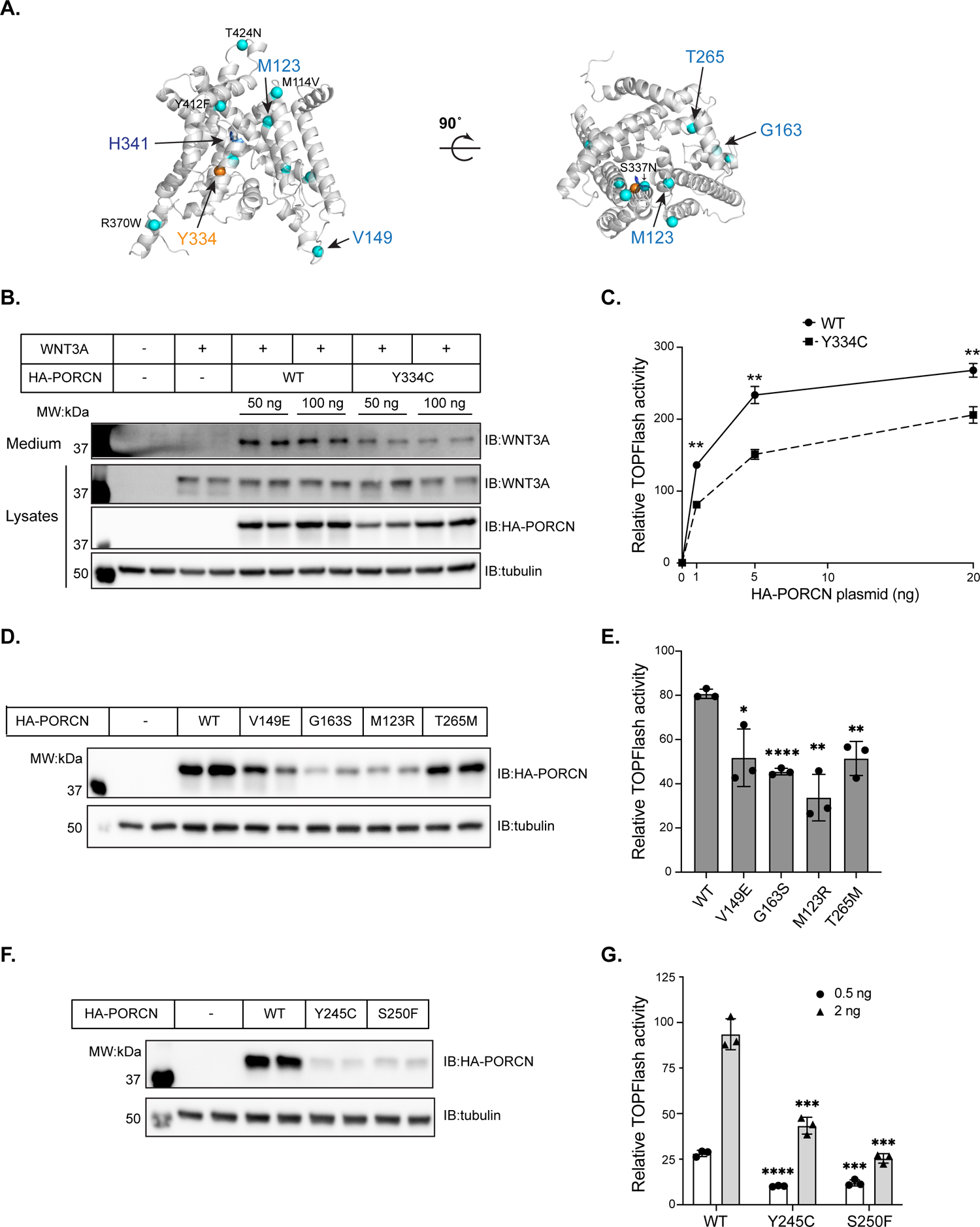
PORCN structure aids in prediction of mutant stability and function. **(A).** PORCN VUS mutants M114V, S337N, R370W, T424N and Y412F are labeled on the predicted PORCN homology model by blue spheres. The mutants described in the text (M123R, V149E, G163S and T265M) are highlighted by arrow heads. The catalytic His341 is shown as blue sticks. Y334 is colored orange. **(B).** The PORCN Y334C mutant has decreased protein abundance in lysates, leading to decreased WNT3A secretion into culture medium. HT1080 PORCN KO cells were transfected with WNT3A and the indicated ng of PORCN expression plasmids. PORCN expression in lysates and WNT3A secretion into the medium were assessed by immunoblotting. **(C).** The Y334C mutant has decreased activity in a Wnt/β-catenin signaling assay, consistent with decreased protein expression. HT1080 PORCN KO cells were transfected with TOPFlash reporter, as well as mCherry, WNT3A and the indicated PORCN expression plasmids. The significance of the difference was assessed by unpaired t test comparing each data point of Y334C activity with WT plasmids. Error bars represent standard deviation. **, P<0.01. **(D).** Several PORCN VUS mutants have reduced expression levels. HT1080 PORCN KO cells were transfected with 50 ng of the indicated PORCN expression plasmids and expression was assessed by immunoblotting of cell lysates. **(E).** Predicted deleterious PORCN VUS mutants have reduced activity in the Wnt reporter assay. Similar to (C), HT1080 PORCN KO cells were transfected with TOPFlash reporter, as well as mCherry, WNT3A and respective PORCN expression plasmids (1.5 ng). The significance of the difference was assessed by unpaired t test comparing each mutant activity with WT plasmids. Error bars represent standard deviation. *, p<0.05; **, p<0.01; ****, p<0.0001. **(F).** Additional PORCN mutants have reduced expression levels. Method as above. **(G).** PORCN mutations found in non-Goltz syndrome male patients have reduced activity in Wnt reporter assay. Similar to (C), HT1080 PORCN KO cells were transfected with TOPFlash reporter together with mCherry, WNT3A and PORCN expression plasmids.

### Characterization of additional PORCN VUS mutants

From the Global Variome shared LOVD database, we identified 9 PORCN missense VUSs. These variants fall in diverse regions of the predicted protein structure (Figure 4A). We examined the expression and activity of four of the VUSs including V149E, G163S, M123R and T265M that have not previously been biochemically characterized. G163S appeared to be unstable (Figure 4D). Glycine in TM loops has been reported to facilitate helix packing (Javadpour et al., 1999), and indeed Gly163 lies on intracellular loop 1 where the opposite face packs tightly with TM helices 2 and 4 (Supplemental Figure 4B). M123R was also unstable (Figure 4D). Met123 lies on the ER lumenal end of TM helix 4 interacting with two methionines (Met109 and Met112) on TM 3 potentially forming a methionine triad, a structure previously described at the ends of TM loops in the copper transporter CTR1 (Ren et al., 2019) (Supplemental Figure 4C). T265M, which decreases PORCN enzymatic activity without a change in protein stability, has potential to disrupt the binding of PAM-CoA to PORCN as it lies at the end of tunnel 2, adjacent to the end of the fatty acid chain (Supplemental Figure 4D). Both M123R and T265M are predicted as possibly or probably damaging by both PolyPhen-2 (Kelley et al., 2015) and Missense 3.0. V149E, which also has decreased expression, was not predicted by PolyPhen-2 and Missense 3.0 to have deleterious consequences for this mutant. Val149E replaces a conserved hydrophobic residue with a charged residue in the cytoplasmic domain, distant from the catalytic core (Figure 4A). We speculate that this impairs protein stability due to altered protein-protein interactions of the cytoplasmic loop. Consistent with the model prediction as well as the observed protein abundance, all the mutants have reduced Wnt/ý-catenin signaling activity in the reporter assay (Figure 4E).

Goltz syndrome is an X-linked dominant disorder, and it mainly occurs in females. However, male patients with PORCN mutations who do not display typical FDH symptoms have been reported (Brady et al., 2015; Madan et al., 2017; Wawrocka et al., 2021). In particular, one family of brothers (hemizygous Y245C) had anophthalmia/microphthalmia (Wawrocka et al., 2021), which is not typically seen in FDH patients. We analyzed two such mutations, Y245C (possibly damaging) and S250F (probably damaging) by Polyphen-2 (Madan et al., 2017). However, both mutants have very low expression levels as assessed by western blotting (Figure 4F), and thus reduced reporter activity (Figure 4G). Tyr245 on TM helix 6 packs tightly with Leu214 and Phe218 on TM5 (Supplemental Figure 4E); its replacement by Cys may disrupt this network. S250F is predicted to be probably damaging. It lies in the catalytic core, and the serine is part of a network with His409 of TM9 and Wnt hairpin 2 (Supplemental Figure 4F). Replacement by the more bulky and hydrophobic phenylalanine may disrupt this network. Again, the structure of PORCN assists in understanding the structural consequences of patient-associated variants.

### Docking ETC-159 and other PORCN inhibitors illustrates drug binding modes

Pharmacologic inhibition of PORCN may be efficacious in Wnt-high disorders such as cancer, and a number of candidates have shown robust activity in various model systems (Zhong and Virshup, 2020). We examined if the model could elucidate the binding modes of potent small molecule inhibitors of PORCN. To this end, we used the standard precision mode of GLIDE (Friesner et al., 2006) to dock the PORCN inhibitors ETC-159 (Madan et al., 2016), Wnt-C59 (Proffitt et al., 2013), LGK-974 (also called WNT974) {Liu.2013r}, IWP2 and its derivative IWP-L6 (Wang et al., 2013) as well as ETC-131 in the homology model (see Table 1 for chemical structures and IC_50_ of the drugs). Notably, all the active inhibitors adopt an inverted “L” shape and occupy the same binding site within the TM core, partially overlapping with the PAM-CoA binding (Figure 5A and 5B, Supplemental Figure 5A-E). The biaryl moiety (colored red in Table 1) of the inhibitors sits deeply in the binding cavity (Figure 5B, 5C). In Wnt-C59, LGK-974, and ETC-159, the aromatic ring at the bottom is surrounded by several hydrophobic residues including Val302, Val350, Leu405 and notable, Tyr334, (Figure 5C, and supplemental Figure 5A, 5B).

**Figure 5.**
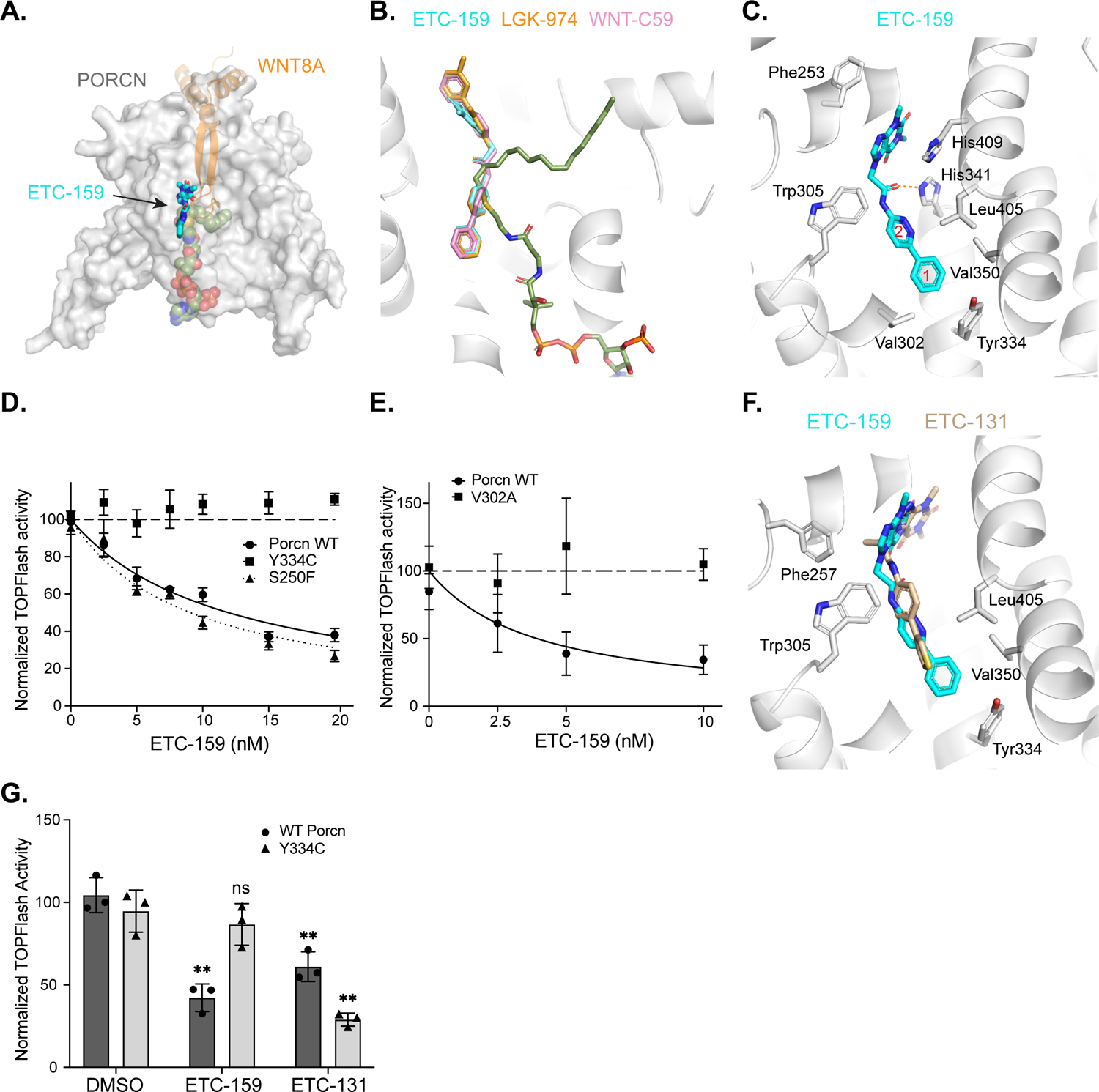
PORCN model predicts mechanisms of Wnt inhibitor binding. **(A).** PORCN inhibitor ETC-159 docked onto PORCN and overlaid on WNT8A and PAM-CoA. The inhibitor occupies the active site, conflicting with both Wnt hairpin 2 and PAM-CoA. **(B).** A close-up view of (A) showing the predicted binding modes of ETC-159, LGK-974 and WNT-C59, and highlighting the conflict with binding of palmitoleoyl-CoA. **(C).** A close-up view of ETC-159 docking as determined by Protein-Ligand Interaction Profiler (PLIP) (Adasme et al., 2021) predicts hydrophobic interactions with PORCN Leu405 and Val350 as well as the Val302 and the disease-associated residue Tyr334. **(D).** PORCN Tyr334C mutant is resistant to inhibition by ETC-159. HT1080 PORCN KO cells were transfected with TOPFlash reporter, and mCherry, WNT3A and respective PORCN expression plasmids. 1 ng of WT PORCN, 2 ng of Y334C and 5 ng of S250F mutant plasmids were used in each assay to get approximately similar basal activities. ETC-159 was added 6 hours after transfection and cells were harvested 24 hours later. **(E).** PORCN V302A mutant is resistant to inhibition by ETC-159. Similar to Figure 5D, except 5 ng of V302A mutant plasmids were used. **(F).** A close-up view of ETC-131 (gold) binding mode in PORCN superimposed on ETC-159 (blue) binding, highlighting the greater distance between ETC-131 and PORCN Tyr334. **(G).** Y334C mutant is resistant to inhibition by ETC-159, but not by ETC-131. HT1080 PORCN KO cells were transfected with TOPFlash reporter, and mCherry, WNT3A and respective PORCN expression plasmids. 1 ng of WT and 2 ng of Y334C PORCN expression plasmids were used in each assay to get approximately similar basal activities. 20 nM of ETC-159 or 0.5 nM of ETC-131 was added 6 hours after transfection and cells were harvested 24 hours later. The experiment was repeated three times, with a representative experiment shown. The significance of the difference was assessed by unpaired t test for each plasmid comparing drug treatment with the DMSO control. Error bars represent standard deviation. **, p<0.01.

**Table 1.**
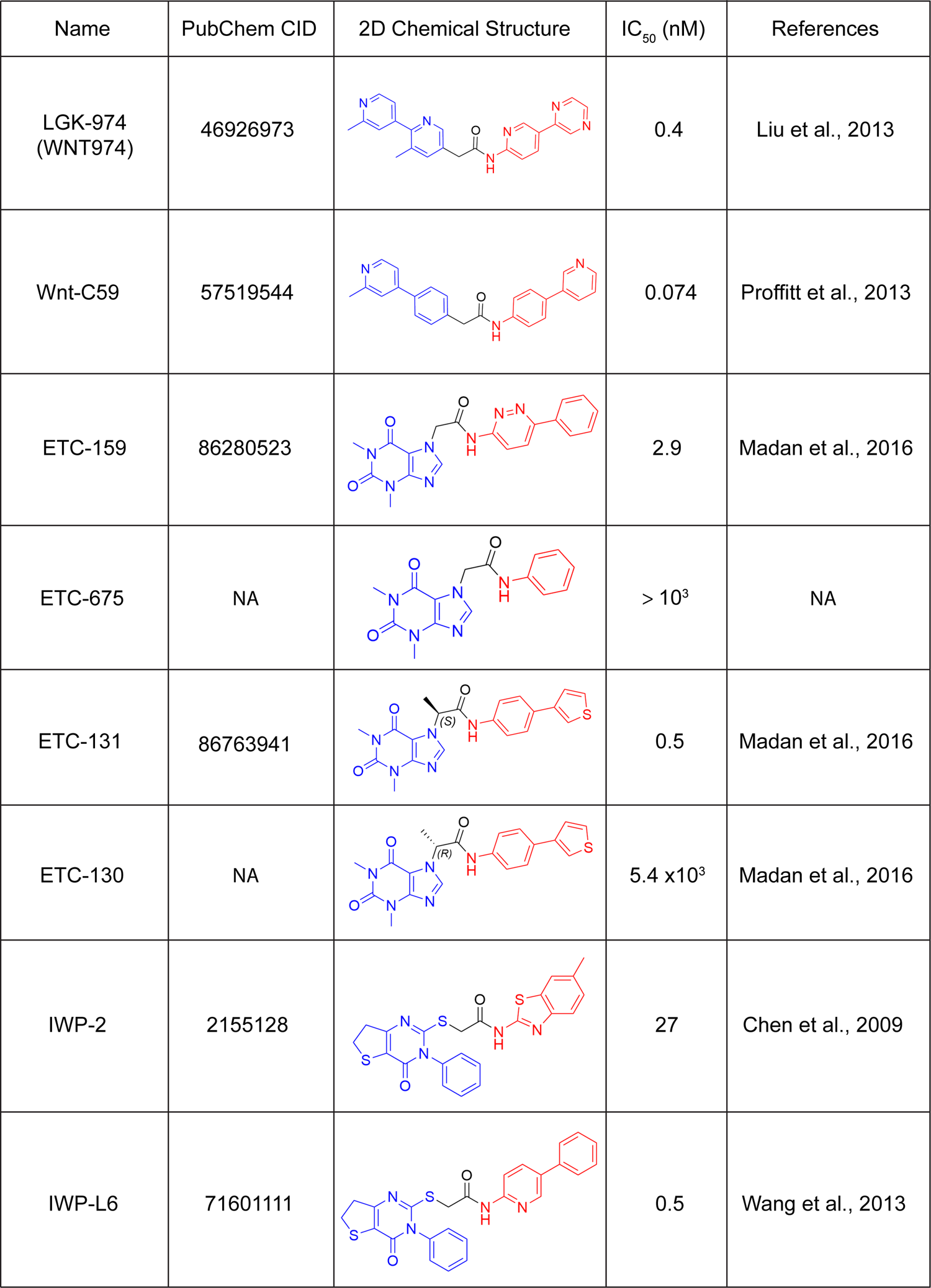
PORCN inhibitors chemical structures and IC50.

| Name | PubChem CID | 2D Chemical Structure | IC<sub>50</sub> (nM) | References |
| --- | --- | --- | --- | --- |
| LGK-974<br>(WNT974) | 46926973 |  | 0.4 | Liu et al., 2013 |
| Wnt-C59 | 57519544 |  | 0.074 | Proffitt et al., 2013 |
| ETC-159 | 86280523 |  | 2.9 | Madan et al., 2016 |
| ETC-675 | NA |  | > 10<sup>3</sup> | NA |
| ETC-131 | 86763941 |  | 0.5 | Madan et al., 2016 |
| ETC-130 | NA |  | 5.4 x10<sup>3</sup> | Madan et al., 2016 |
| IWP-2 | 2155128 |  | 27 | Chen et al., 2009 |
| IWP-L6 | 71601111 |  | 0.5 | Wang et al., 2013 |

We tested the predictions of the model by mutating several of the residues involved in close contacts with inhibitors. Remarkably, and strongly supporting the accuracy of the model, mutating the Val302 to Ala, or Tyr334 to either Cys (i.e. the patient mutation) or Ala made PORCN resistant to inhibition by ETC-159 (Figure 5D, 5E, Supplemental Figure 5G). This is not a general feature of disease-associated variants, as Ser250 mutated to Phe, found in two males with FDH, did not confer drug resistance (Figure 5D) (Madan et al., 2017). Previous structure-activity relationship studies indicated that ETC-675, a close analogue of ETC-159 that does not have the bottom aromatic ring, completely lost its inhibitory effect (IC_50_ > 1 μM) in the TOPFlash assay (Table 1). These results, taken together, suggest that the interactions of the aromatic ring with residues including Tyr334 and Val302 at the bottom of the binding site are indispensable for the inhibitors’ activity.

Towards the ER lumenal side of the catalytic pocket, the second aromatic ring (ring 2 in Figure 5C; colored red in Table 1) of the active inhibitors is wedged in between Trp305 and His341 (Figure 5C, 5F and Supplemental Figure 5A, 5B, 5F). The carbonyl oxygen of the amide linker was predicted to form a hydrogen bond with the imidazole NH of His341 (Poulsen et al., 2015). The cap xanthine ring of ETC-159 is sandwiched between Phe253 and His409 and forms π-stacking interactions with these two residues (Figure 5C). Overall, the superimposed alignment of the predicted inhibitor binding modes agrees with a previously reported ligand-based pharmacophore model of the PORCN inhibitors (Poulsen et al., 2015). In contrast, our model predicted that ETC-131, IWP-2, and IWP-L6 bind at a position which is slightly higher as compared to the docking site of the other inhibitors (Figure 5F and Supplemental Figure 5D-5F). If the binding mode is accurate, the bottom thiophene ring of ETC-131 will be further away from PORCN Tyr334 than the aromatic ring #1 of ETC-159. In other words, mutation of Tyr334 should have less effects on drug sensitivity for ETC-131 than ETC-159. We therefore tested this experimentally. As shown in Figure 5G, although Y334C is completely resistant to ETC-159, it is still sensitive to ETC-131, supporting the prediction model. ETC-131 has a methyl group (S-isomer) attached to the carbon atom between the amide linker and the xanthine ring. Based on the binding mode of ETC-159, we observed that an extra methyl is difficult to accommodate due to the presence of two bulky residues, Phe257 and Trp305, next to the carbon atom between the amide linker and the xanthine ring. These findings may explain the slight difference of the binding mode as adopted by ETC-131 (Figure 5F and Supplemental Figure 5D). More interestingly, molecular docking of ETC-130 (the R-isomer of ETC-131) positioned the ligand in the binding site with an upside-down flip orientation (Supplemental Figure 5C), which agrees with the experimental result showing that ETC-130 is inactive in the TOPFlash assay (Madan et al., 2016). Despite containing a biaryl moiety that can insert deeply into the binding cavity of PORCN, IWP-L6 (a derivative of IWP-2) (Chen et al., 2009; Wang et al., 2013) has a much bulkier group at the lumenal side of the protein binding tunnel. Not surprisingly, the docking results positioned IWP-L6 in a similar binding location relative to ETC-131 (Supplemental Figure 5F). Collectively, the modeling results strongly indicate that the model can provide useful insights and predictions into the molecular interactions between PORCN and its inhibitors.

## DISCUSSION

PORCN is the sole acyltransferase responsible for the palmitoleation of all Wnts. It is a promising drug target for Wnt-addicted cancers, and several PORCN inhibitors are currently in clinical trials. Loss-of-function mutations in PORCN cause FDH predominantly in females due to loss of Wnt signaling during development. However, X chromosome inactivation as well the large number of alleles cause a wide range of manifestations of FDH, making both the clinical diagnosis and the classification of new variants of uncertain significance (VUS) challenging. Here, building on recent structures of other human MBOAT proteins and as part of an effort to characterize a VUS in PORCN, we developed a structural homology model. The usefulness of the model is supported by its ability to predict the docking of WNT8A and PAM-CoA, reveal the consequences of the patient’s Y334C mutation, assist in understanding and classifying other VUSs, and finally, understanding the structure-activity relationship of multiple small molecule inhibitors of PORCN.

The PORCN model was templated on the recently published DGAT1 cryo-EM structures. DGAT1 catalyzes the transfer of the acyl group from oleoyl-CoA (*cis*-C18:1) to cholesterol to form cholesterol esters. Like PORCN, DGAT1 prefers unsaturated fatty acids over saturated acyl groups. (Asciolla et al., 2017; Chang et al., 2010; Rios-Esteves and Resh, 2013; Tuladhar et al., 2019). The accuracy of the PORCN model is demonstrated by our ability to dock palmitoleoyl-but not saturated palmitoyl-CoA into the PORCN tunnel 2. We note that the kinked acyl chain tunnel was also observed in the structures of stearoyl CoA desaturase (SCD), the enzyme that desaturates stearoyl and palmitoyl-CoA, and Notum, an esterase that cleaves the palmitoleoyl moiety from Wnts (Bai et al., 2015; Kakugawa et al., 2015). In PORCN, tunnel 2 accommodating the palmitoleoyl chain is surrounded by both highly conserved and co-evolving residues (Figure 3E), suggesting this recognition of a mono-unsaturated substrate is important throughout the animal kingdom.

Identifying a correct and optimal template for building a 3D model backbone is the first and the most critical step in the process of homology modelling. Templates for modeling PORCN were initially selected by automated homology modeling programs like Phyre2 based on sequence alignment. The bacterial MBOAT protein DltB was first used due to slightly higher identity compared to other human MBOAT (14% for DltB and 13% for DGAT1 in Phyre2). However, the tunnels in DltB that accommodates small D-alanyl groups, are very different from those in the human MBOAT proteins (Supplemental Figure 3A). Furthermore, DltB is on the bacterial plasma membrane while PORCN, ACAT1 and DGAT1 are located on the mammalian ER membrane, two membranes with very different compositions (Holthuis and Menon, 2014). Thus, the homology model based on DltB appeared less predictive than that based on DGAT1.

The predicted structure also suggests a pathway for Wnts to be acylated by PORCN and then handed off to the carrier protein WLS in the ER membrane. In the model, Wnt hairpin 2 enters tunnel 1 from the lumenal face of the ER. Wnt hairpins 1 and 3 make additional contacts with the lumenal loops of PORCN to stabilize the interaction. Following PAM-COA docking, the catalytic His341 activates WNT8A Ser186 for nucleophilic attack on the thioester bond. The next step, the transfer of acylated Wnt from PORCN to WLS, is probably direct (Yu et al., 2014). In both the WNT8A: PORCN complex (here) and the WNT8A: WLS complex (Nygaard et al., 2021) Wnt hairpin 2 and the attached palmitoleoyl-group are inserted into the transmembrane domain. This suggests that the most energetically favorable route for acylated Wnt is a lateral transfer of the hairpin 2-PAM from PORCN to WLS within the plane of the membrane. Inspecting the ternary PORCN: WNT8A: PAM-CoA complex, we speculate that the most likely route for Wnt transfer is to slide between PORCN TM helices 3 and 4 on one side, and TM10 on the other. Consistent with this mode of transfer, we note that the conformation of WNT8A used here was essentially unchanged from its conformation bound to WLS (Nygaard et al., 2021), and that WNT3 in its FZD CRD bound conformation (PDB 6AHY) could not dock to PORCN in a reasonable position (Hirai et al., 2019). This prediction of WNT lateral transfer from PORCN to WLS may be testable in future studies.

While the current PORCN model has demonstrated its applicability in providing an overall framework of both Wnt palmitoleation and inhibitor binding, it has several caveats that warrant further cautions. First, the protein-protein docking model was not able to predict the bona fide binding geometry between the catalytic His341 and the Ser186 on the hairpin loop 2 of Wnt-8A, despite a distance of 6.3 Å between the NH of the imidazole from His341 and the hydroxyl group of Ser186 from Wnt-8A. The model did not fully explain the role of Asn306, which has been proposed to aid catalysis but whose mutation did not affect function in two cell-based assays. Finally, our in-house structure–activity relationship (SAR) analysis and a previous pharmacophore model of PORCN inhibitors indicated that the oxygen atom from the xanthine ring plays a role as a hydrogen bond acceptor during protein-ligand interaction (Poulsen et al., 2015). Our model failed to identify the exact protein residue that forms the hydrogen bond with the oxygen from the xanthine group.

These uncertainties may arise from the availability of limited choice of template for the modeling, as well as the low sequence homology between the localized protein motif. However, it is also possible that the pharmacophore model was too simplistic, assuming that the biphenyl substituents of all inhibitors are perfectly superimposable in the active site. Atomic resolution structures of MBOAT protein family members have only been available recently. Future work focusing on extensive model refinement with the availability of new structures deposited in the public protein data bank, together with the aid of site-directed mutagenesis should facilitate the improvement of the model. Meanwhile, we believe an experimentally determined three-dimensional structure of the Wnt-PORCN complex or an inhibitor-bound PORCN structure will address the uncertainties presented herein.

## ACKNOWLEDGEMENTS

The authors thank members of the Virshup lab, including Babita Madan for discussions on PORCN structure and potential enzymatic mechanisms, and Yunka Wong for technical assistance. We had invaluable assistance with LERI from Ngaam J. Cheung (aka Yaan), University of Oxford).

