## Supplemental Figures 1-5 for "Structural model of PORCN illuminates disease-associated variants and drug binding sites"

#### Supplemental Figure 1. Multiple sequence alignment of diverse PORCN with other MBOAT family members.

Sequences of PORCN from human (NP_982301.1), mouse (NP_076127.1), *C.elegans* (NP_509054.1) and *Drosophila* (NP_476890.1) as well as human ACAT1 (NP_003092.4), DGAT1 (NP_036211.2) and GOAT (Ghrelin O-acyltransferase, or MBOAT4, NP_001094386.1) were obtained from Uniprot. They were aligned using the Clustal Omega online server (https://www.ebi.ac.uk/Tools/msa/clustalo/) and colored using Jalview with the ClustalX color scheme. The catalytic Histidine and conserved Asparagine are outlined in the orange boxes, and the mutant human Y334 is indicated with a red arrow.

### Supplemental Figure 1 .

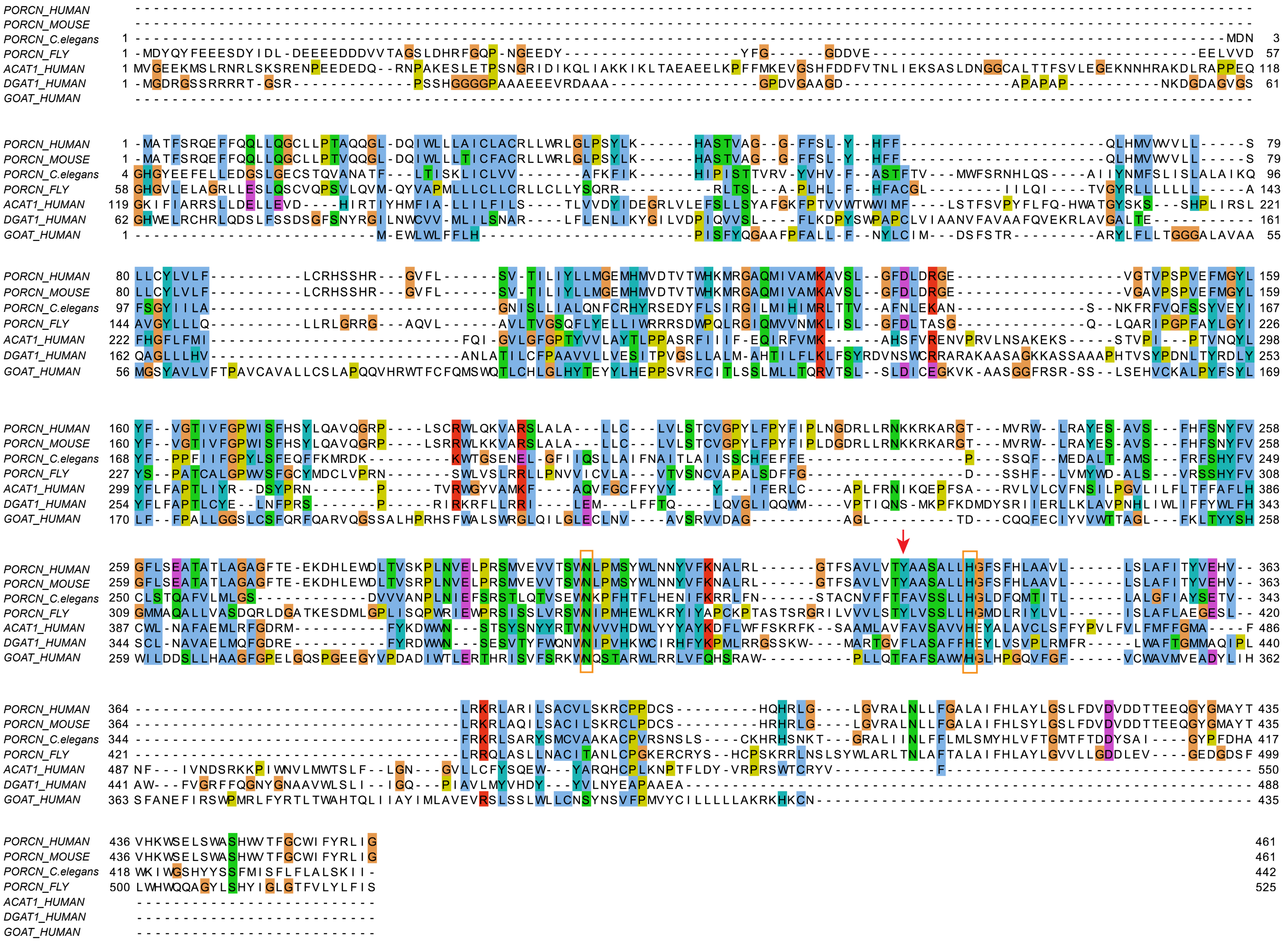

#### Supplemental Figure 2. Additional features of the PORCN homology model.

**(A).** Alignment of the predicted PORCN structure with ACAT1 (PDB 6P2P) (left, wheat color) and DltB (PDB 6BUH) (right, pale green color). ACAT1 has a root mean square deviation (RMSD) of 5.3 Å and DltB has an RMSD of 5.2 Å with PORCN (Cα backbone).

**(B).** Conservation of PORCN model in surface presentation, three views. Conserved residues cluster in the catalytic core.

**(C).** Co-evolving PORCN residues, with the strongest coupling block outlined in blue. From the Leri website with the assistance of Ngaam J. Cheung.

**(D).** Example of co-evolution of the PORCN catalytic core. Selected image from the multiple sequence alignment of 468 PORCN sequences, with amino acids represented in color as follow: small nonpolar: G, A, S, T; hydrophobic: C, V, I, L, P, F, Y, M, W; polar: N, Q, H; negatively charged: D, E; and positively charged: K, R. The boxed residues have the highest coupling scores, i.e., fall in the bottom left corner of Supplemental Figure 2C. The nearly invariant catalytic histidine 341 is indicated with a red star.

Supplemental Figure 2 .

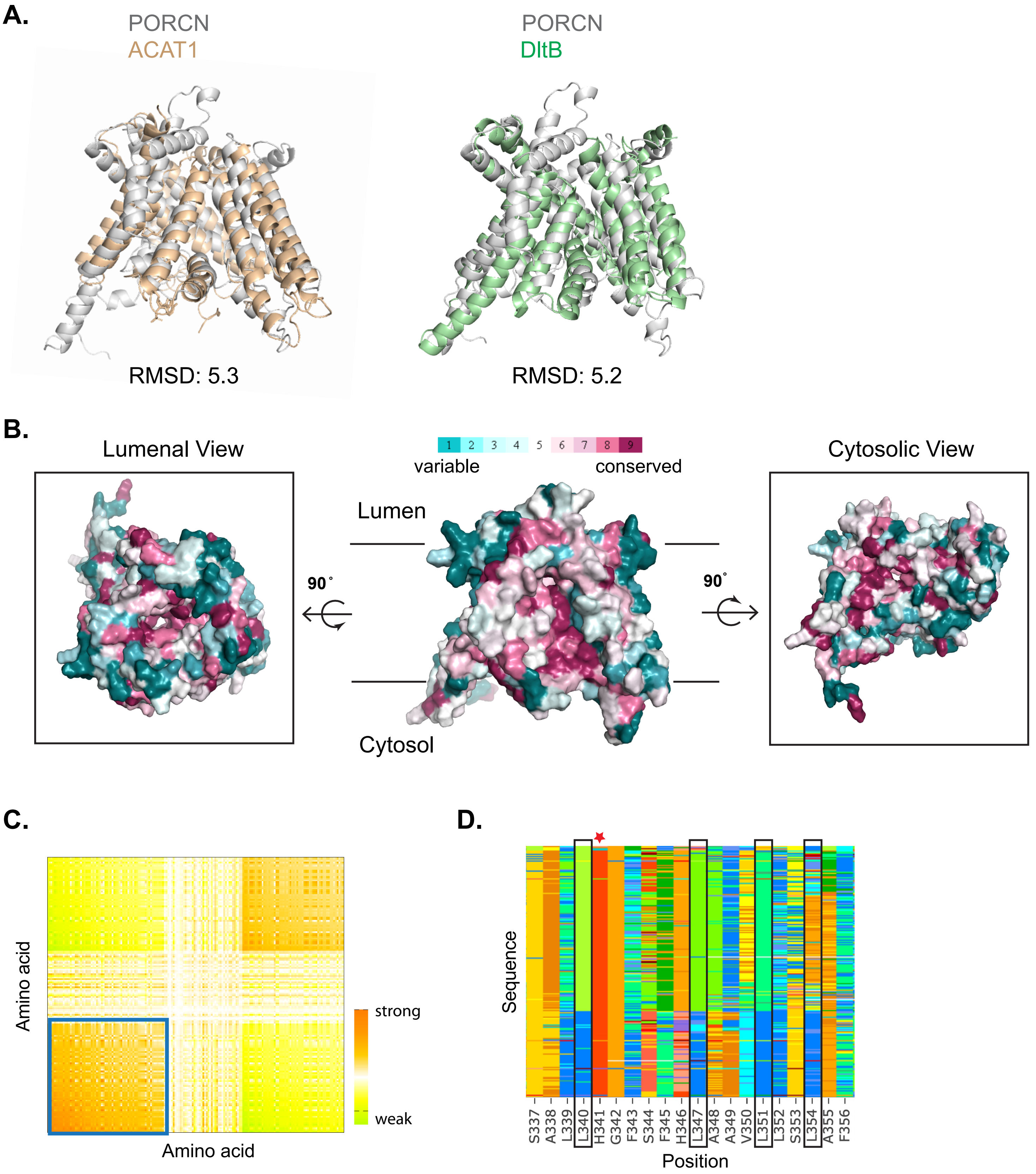

#### Supplemental Figure 3. PAM-CoA and WNT8A docking into predicted PORCN model

**(A).** The tunnels in the indicated MBOAT structures are illustrated. DltB (PDB 6BUH), DGAT1 (PDB 6VP0), HHAT (PDB 7MHY).

**(B).** Workflow for generating the PORCN-WNT8A-PAM CoA complex. PAM-CoA was first docked into the PORCN tunnel 2 (step 1). Independently, human WNT8A (from PDB 7KC4) was docked into the PORCN lumenal tunnel 1 (step 2). Finally, the two docking results were superimposed to show the complex structure.

**(C).** PORCN tunnel 2 is lined with hydrophobic residues. A sliced surface view of PORCN is shown to highlight the hydrophobic cavity that is occupied by the acyl chain of PAM-CoA. PAM-CoA is represented by green sticks.

**(D).** A close-up view of the residues that are close to the *cis*-double bond of acyl chain of PAM-CoA, including the conserved Asn306 and Met309 on PORCN and Ser190 on WNT8A. PAM-CoA is represented by green sticks.

**(E).** PORCN-PAM CoA model predicts the deleterious effects of mutants tested by Resh and co-workers {Rios-Esteves.2014}, highlighting Trp305, Tyr334, Ser337 and Leu340 on PORCN. PAM-CoA is represented by green sticks.

**Supplemental Figure 3.**

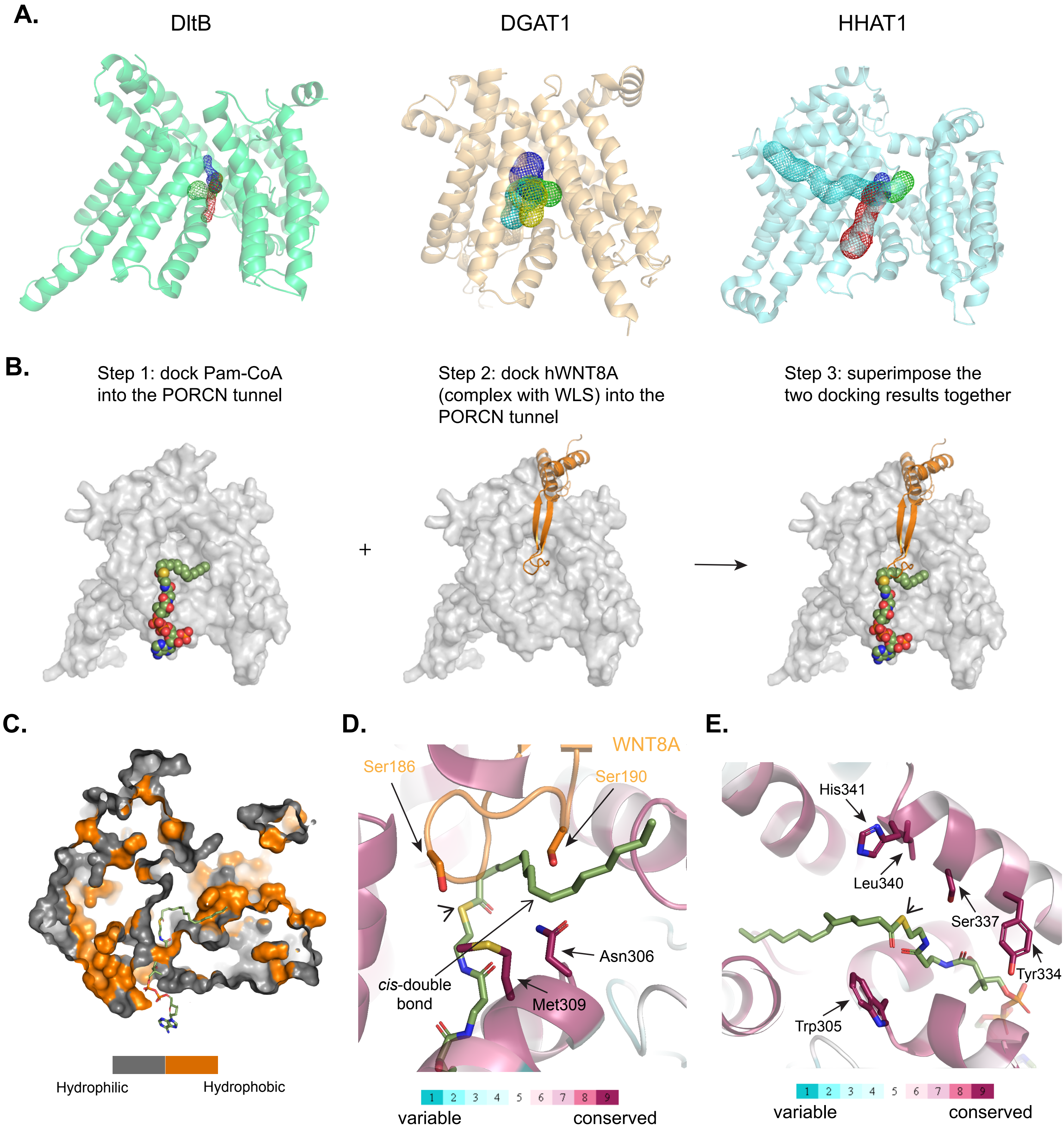

#### Supplemental Figure 4. Patient variants are predicted to disrupt PORCN structure.

**(A).** Tyr334 (Y334C in a patient) is located on TM7 and interacts with neighboring hydrophobic residues such as Val302 and Leu354.

**(B).** Gly163 (G163S in a patient) on IL1 is closed packed to TM2 and TM4.

**(C).** Met123 (M123R in a patient) is located on TM4 and forms a methionine triad with Met112 and Met109 on TM3.

**(D).** Thr265 (T265M in a patient) is at the end of the acyl chain of PAM-CoA.

**(E).** Tyr245 (Y245C in a patient) is located on TM6 and interacts with Leu214 and Phe218 on TM5.

**(F).** Ser250 (S250F in a patient) is positioned in the catalytical core together with His409 and His341.

**Supplemental Figure 4.**

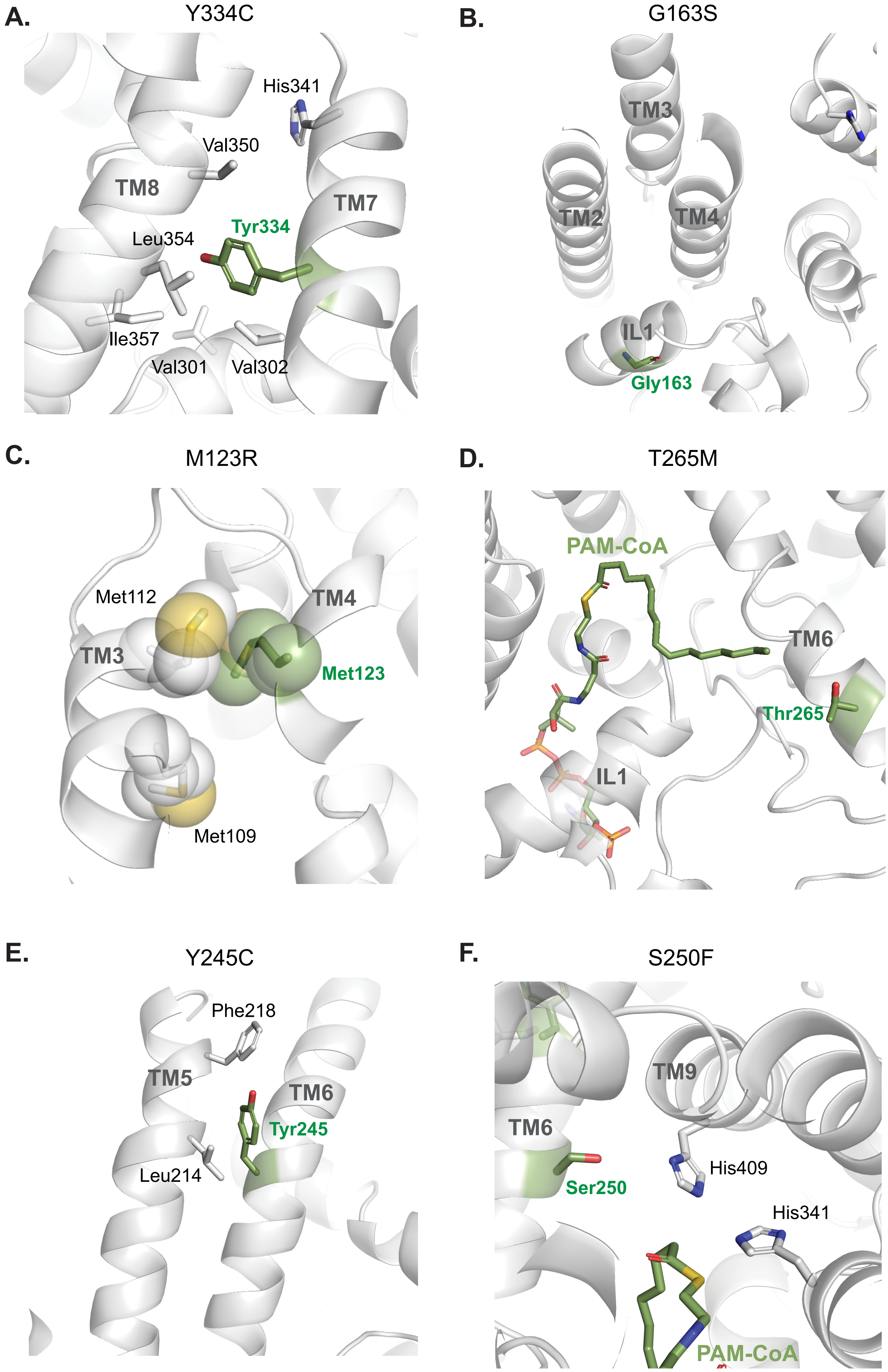

#### Supplemental Figure 5. Wnt inhibitors dock into the PORCN model

**(A-F).** A close-up view showing the predicted binding modes of LGK-974, WNT-C59, ETC-130, ETC-131, IWP2, and IWP-L6 respectively, highlighting the importance of the bi-phenol group at the bottom of LGK-974 and WNT-C59 that could interact with Y334 of PORCN.

**(G).** Both Y334C and Y334A are resistant to inhibition by ETC-159. Similar to Figure 5D, HT1080 PORCN KO cells were transfected with TOPFlash reporter, and mCherry, WNT3A and respective PORCN expression plasmids. 1 ng of WT PORCN plasmids, 2 ng of Y334C, 10 ng of Y334A and 5 ng of L405A mutant plasmids were used in each assay to get approximately similar basal activities without drug treatment. 20 nM of ETC-159 was added 6 hours after transfection and cells were harvested 24 hours later. The relative TOPFlash activity was normalized to each PORCN plasmid under DMSO condition.

**Supplemental Figure 5.**

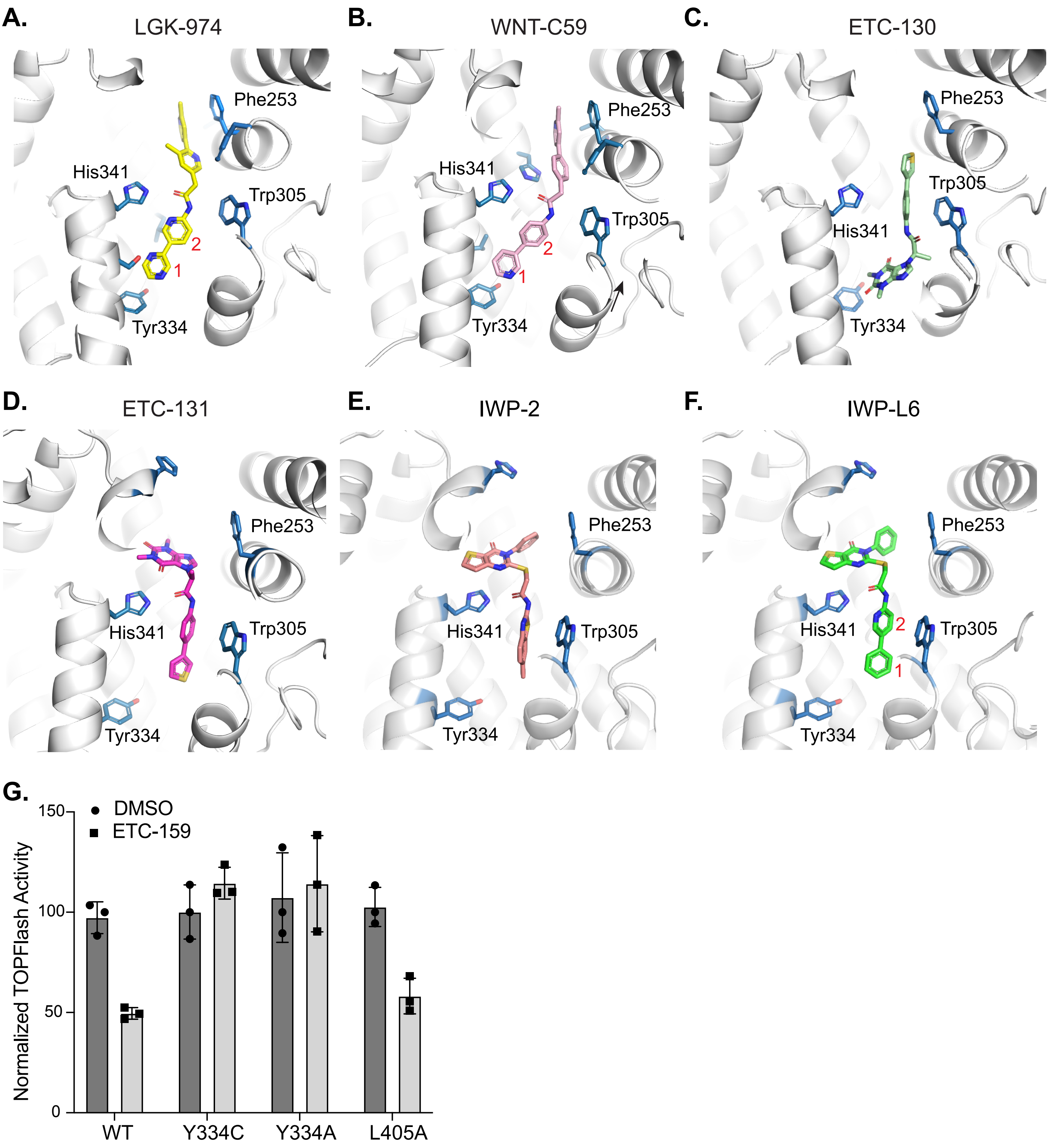
